# First-in-class matrix anti-assembly peptide prevents staphylococcal biofilm *in vitro* and *in vivo*

**DOI:** 10.1101/2020.04.03.022020

**Authors:** Rafael Gomes Von Borowski, Sophie Chat, Rafael Schneider, Sylvie Nonin-Lecomte, Serge Bouaziz, Emmanuel Giudice, Aline Rigon Zimmer, Simone Cristina Baggio Gnoatto, Alexandre José Macedo, Reynald Gillet

## Abstract

Staphylococci are pathogenic biofilm-forming bacteria, source of multidrug-resistance and/or – tolerance causing a broad spectrum of infections. These bacteria are enclosed in a matrix that allows them to colonize medical devices such as catheters and tissue, and which protects against antibiotics and immune systems. Advances in antibiofilm strategies for targeting this matrix are therefore extremely relevant. Plants are constantly attacked by a wide range of pathogens, and have protective factors such as peptides to defend themselves. These peptides are common components in *Capsicum* peppers (CP). Here, we describe the development of CP bioinspired peptide “capsicumicine”. We demonstrate that capsicumicine strongly prevents methicillin-resistant *S. epidermidis* biofilm *via* a new extracellular “matrix anti-assembly” mechanism of action. Catheters pre-coated with capsicumicine decreased *S. aureus* colonization leading to the attenuation of infection, decreasing mice systemic infection. Capsicumicine is the first-in-class non-antibiotic, carbohydrate-binding peptide.

## Introduction

Antimicrobial failure is a worldwide challenge, currently addressed by a WHO global action plan (*1*). A lack of new antibiotics and the inappropriate use of older treatments mean that multidrug-resistant strains are increasing (*2*). This process is favored by biofilm development, and microorganisms enclosed in biofilm matrix have antibiotic resistance that is up to 1000 times higher than planktonic ones (*3*). This makes the matrix itself an important target for biofilm control. Biofilms are organized microbial clusters made of a self-assembled matrix that usually attaches to a surface, whether abiotic (medical devices such as catheters, teeth, etc.) or biotic (host tissues, mucus, chronic wounds, etc.) (*4, 5*). Since the bacteria are embedded into this matrix, they are harder to treat because of their increased tolerance and resistance to antibiotics, disinfectants, and host defenses (*6, 7*). Other advantages over planktonic forms include physiological and biochemical changes, beneficial quorum sensing, higher (up to 100 times) mutation rates, and persister cell development (*8-10*).

Staphylococci are the most frequent sources of nosocomial infections, particularly *S. epidermidis* and *S. aureus.* While *S. aureus* expresses many virulence factors such as toxins and proteases, for *S. epidermidis* the formation of biofilm is the most important mechanism in infection development (*11*). *Staphylococcus epidermidis* is the most frequent coagulase-negative staphylococcal (CoNS) infection-causing disease (*12*), surviving on various surfaces for months (*13*). It is present in 30% of health care-involving bloodstream infections, and is significantly associated with medical device infections, including 15-40% of prosthetic valve endocarditis (*14*) and 30-43% of prosthetic orthopedic device infections (*15*). More than 150 million intravascular catheters are used per year in the USA, and there are about 250,000 catheter-related infections (*16, 17*). These bacteria are developing antibiotic multi-resistances such as elevated glycopeptide minimal inhibitory concentrations (*18*), and 73-88% of isolates display resistance to oxacillin, fluoroquinolones, macrolides, clindamycin, and trimethoprim/sulfamethoxazole (*19-21*).

In this circumstance, the extracellular matrix is a complex physicochemical barrier representing one of the biggest challenges in microbial treatments (*4*). Therefore, the development of antivirulence strategies such as antibiofilm agents is crucial for the current antibiotic crisis, and peptides are an increasingly arsenal for controlling pathogenic biofilms (*22-24*).

Here, we describe the discovery of the antibiofilm peptide capsicumicine, inspired by natural peptides from the seeds of the red pepper *Capsicum baccatum*. We elucidated its mechanism of action (MoA), the matrix anti-assembly (MAA). Capsicumicine is the first-in-class non-antibiotic peptide displaying this extracellular MoA. Notably, we report an *in vivo* anti-infective proof of concept towards to the use of capsicumicine for complementary treatment of infectious diseases.

## Results

### Capsicumicine prevents biofilm formation without antibiotic activity

We synthesized three peptides inspired by a natural antibiofilm fraction previously identified from *Capsicum baccatum* var. *pendulum* pepper seeds (*25*): P1 (RVQSEEGEDQISQRE), P2 (RAEAFQTAQALPGLCRI), and P3 (RSCQQQIQQAQQLSSCQQYLKQ). To find the most active one, we exposed these compounds to strong biofilm-forming *S. epidermidis* RP62A (ATCC 35984). After 24 h, crystal violet was used to quantify the remaining biofilm (Fig. 1A). The P3, named “capsicumicine,” was the most active with particularly strong antibiofilm activity. Biofilm decreases were observed at all tested concentrations, but especially at 10 μM. There, biofilm was reduced by over 91%, independently of cell growth inhibition (Fig. 1B). To examine the effects of capsicumicine on growth, we checked *S. epidermidis* colony-forming unit (CFU) counts after peptide exposure. As expected, the CFUs were unchanged by capsicumicine, so the peptide’s biofilm inhibition is not due to bactericidal activity (Fig. 1B). To verify the interactions between capsicumicine and established matrices, we exposed a pre-existing *S. epidermidis* biofilm to a single concentration (100 μM) of the peptide. At that concentration, capsicumicine accounts for 15% of the disruption of pre-existing biofilm (Fig. S1).

**Fig. 1.**
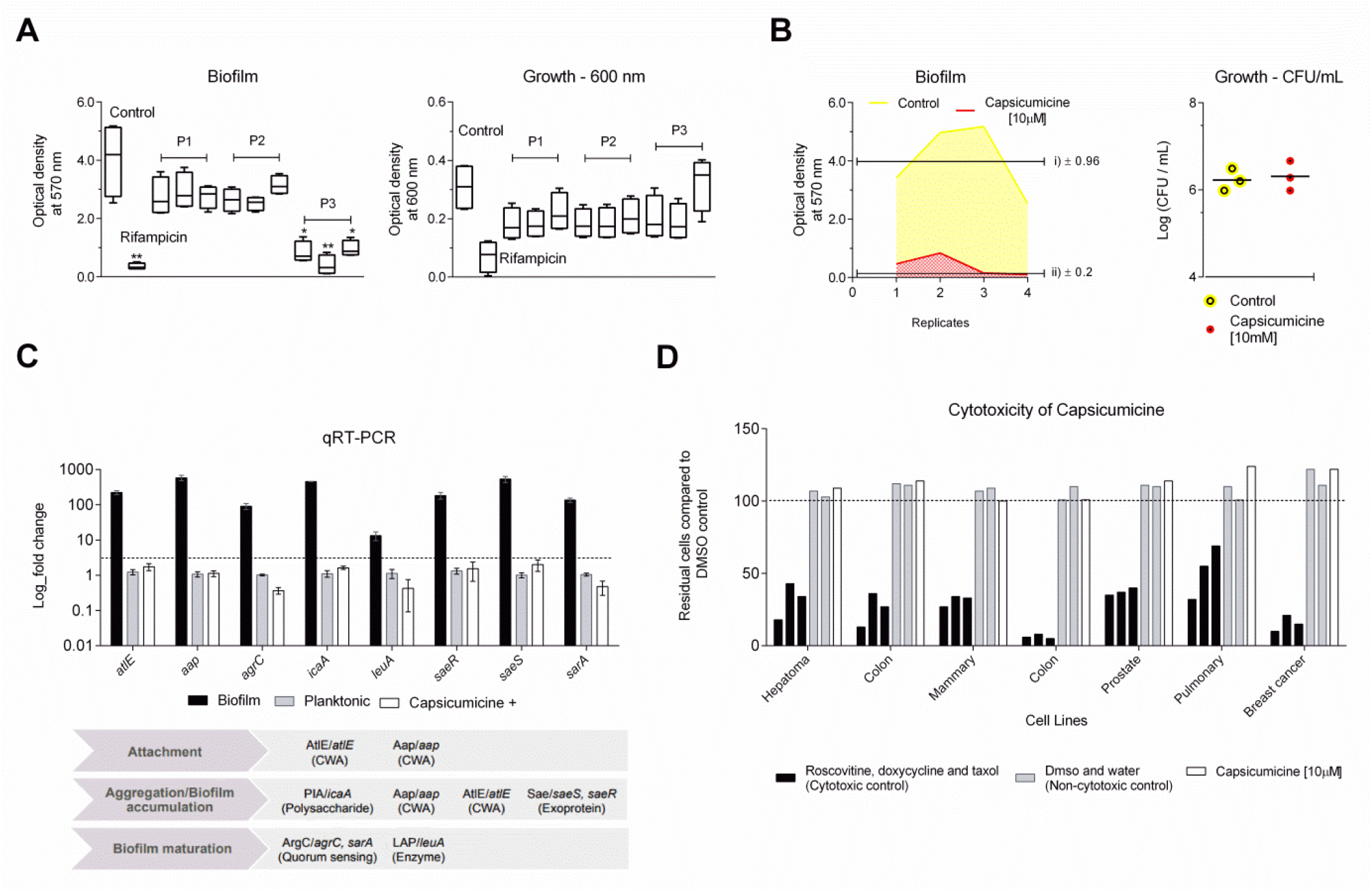
Antibiofilm activity of bioinspired peptides and cytotoxicity. *(A)* Antibiofilm activity of peptides P1, P2, and P3 (capsicumicine) at 1, 10, and 100 μM. Quantification of *Staphylococcus epidermidis* (ATCC 35984) biofilm and growth were done at an optical density; bacteria without peptide exposure (Control) and the antibiotic control, 16 μg/mL rifampicin (Rif.). Student’s *t*-test: *, *p* ≤ 0.05; **, *p* ≤ 0.01. *(B)* Antibiofilm activity and growth of capsicumicine at 10 μM. Yellow represents 100% of formed biofilm and red represents the remaining biofilm after exposure to capsicumicine. Growth was verified using colony-forming units (CFUs). *(C)* Gene expression (mean log fold changes ± standard errors of the means) of the encoding genes involved in *S. epidermidis* biofilm formation as compared to the planktonic (grey) and biofilm controls (black), with the *ssrA* gene used as a reference. The group exposed to capsicumicine is shown in white. CWA, cell wall-anchored proteins. *D*) Capsicumicine cytotoxicity evaluation in representative human cell lines shown via automated image-based cellular content analysis. Cell counts are presented as residual cell percentages (%) compared to the average of the DMSO control, with water control also shown (grey bars). The black bars on the left show cytotoxic controls (roscovitine, doxycycline, and taxol), while the white bars are cells exposed to 10 μM of capsicumicine.

To study the peptide’s possible mechanisms of action, we selected several genes involved in different stages of biofilm development (*atlE, aap, agrC, icaA, leuA, saeR, saeS*, and *sarA*); primers are listed in Table S1. Fold changes were analyzed by quantitative real-time PCR (qRT-PCR). Since exposed bacteria remain planktonic, we compared their relative gene expressions to planktonic control cells. For all tested genes, capsicumicine-exposed cells show the same fold changes as the control (Fig. 1C).

### Capsicumicine is not cytotoxic in mammalian cells

To ensure that capsicumicine is safe before propose *in vivo* trials, we verified its cytotoxicity in seven different representative human cell lines. We used automated image-based cellular content analysis and found that capsicumicine-treated cells perform exactly the same as untreated controls, displaying no cytotoxicity (Fig. 1D).

### Independently of cell interactions, capsicumicine impairs biofilm attachment, aggregation, and accumulation

To explore its activity during the first stages of biofilm development, we analyzed biofilm cultures on polystyrene coupons with and without capsicumicine after 1, 4, and 24 h. Scanning electron microscopy (SEM) analysis shows that bacterial attachment decreases after 1 h of capsicumicine exposure, with biofilm accumulation and cell aggregation profiles strongly reduced after 4 and 24 h (Fig. 2A). This demonstrates that capsicumicine prevents *S. epidermidis* coupon adhesion, and notably this activity still occurs after 24 h incubation. In order to localize the peptide, we used capsicumicine conjugated to fluorescein isothiocyanate (capsicumicine-FITC) and confocal fluorescence microscopy (CFM). Analysis of the CFM images showed that capsicumicine-FITC stays associated with extracellular matrix components, not entering into bacterial cells or the walls or membranes (Fig. 2B).

**Fig. 2.**
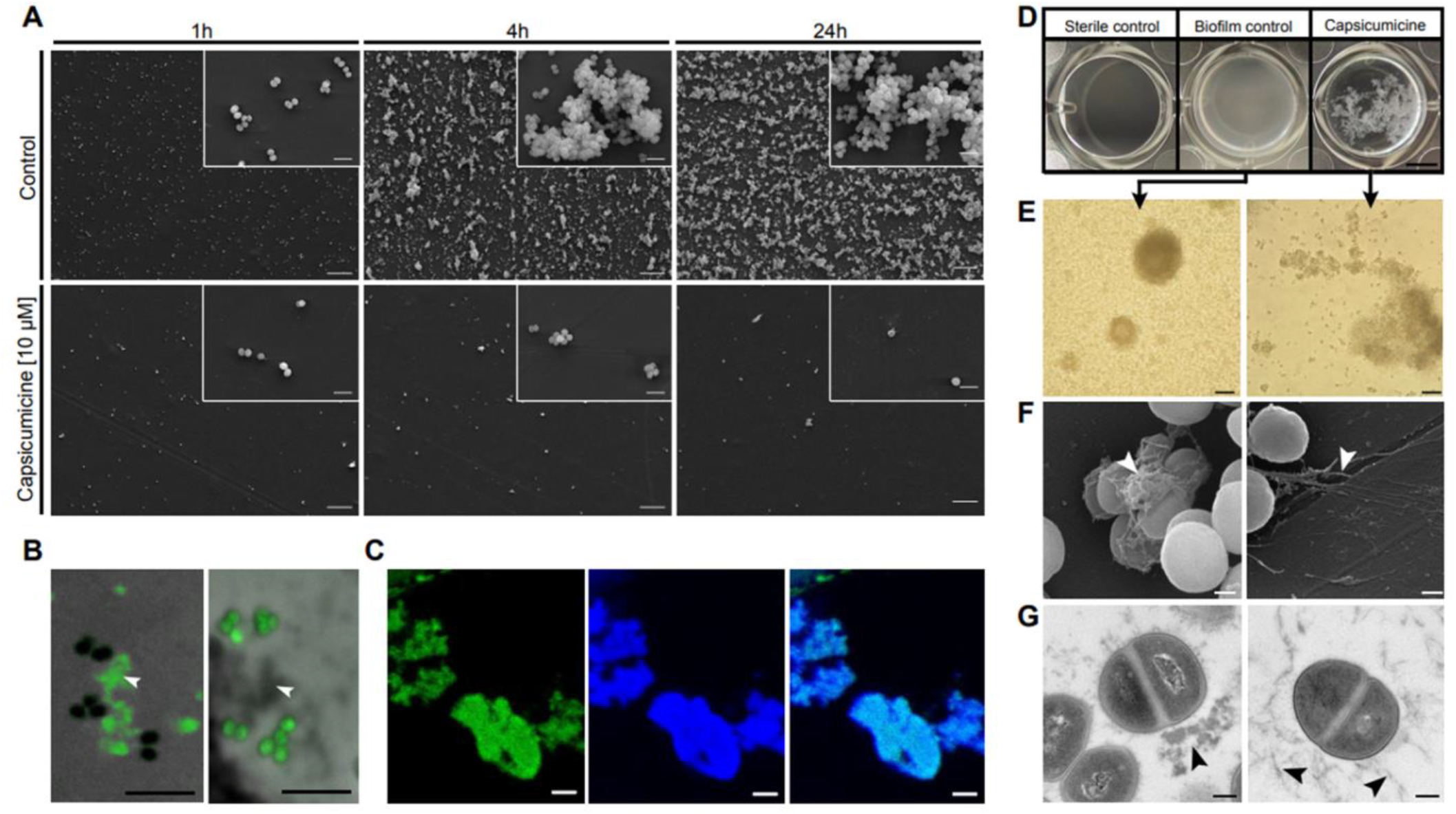
Biological characterization of capsicumicine by microscopic approaches. (*A*) SEM images of polystyrene coupons after 1, 4, or 24h of culture with *Staphylococcus epidermidis* (ATCC 35984). *Top:* peptide-less biofilm control; *bottom:* cultures exposed to 10 μM capsicumicine. Magnification x500, with insets at x5,000; scale bars, 10 μm. (*B*) CFM images: *(left) S. epidermidis* after exposure to 10 μM capsicumicine-FITC, with the matrix in green (fluorescence), and the bacterial cells in black (no fluorescence); (*right*) control, *S. epidermidis* after exposition to pseudonajide-FITC, an antibacterial peptide, this time showing the matrix in gray (no fluorescence) and bacterial cells in green (fluorescence). White arrows indicate matrix content. Scale bars, 5 μm. *(C)* CFM images: *(left) S. epidermidis* after exposure to 10 μM capsicumicine-FITC; *(center)* after exposure to concanavalin A conjugates; and (*right*) their colocalization. Scale bars, 5 μm. *(D-G)* These images explore the organizational state of *S. epidermidis* biofilm after 24 h in the presence (*right*) or absence (control, *left*) of capsicumicine. (*D*) Macroscopic examination by pictures from the bottom of 24-well plates. The “sterile control” shows no bacteria or biofilm formation; the “biofilm control” shows homogenous adhered layers of bacteria; and “capsicumicine” shows non-adhered bacteria but agglutinates; scale bar, 5 mm. *(E)* TLM images show the biofilm control with overlapping attached bacterial clusters, while the capsicumicine-exposed culture shows agglutinated non-adhered cells; scale bar, 20 μm. *(F)* SEM images show the control with dense globular-like matrix features, while the capsicumicine-exposed culture shows fibrillary branch-like oligomer structures; scale bar, 200 nm. (*G*) TEM images of the biofilm control show denser assembled structures, while the capsicumicine-exposed displays thin fibrillary oligomer structures; scale bar, 200 nm. Arrows indicate the matrices.

### Capsicumicine disturbs *S. epidermidis* matrix assembly

Since capsicumicine’s antibiofilm activity was not associated with direct bacterial interactions or gene expression modulation, we used various microscopic approaches to investigate the interactions between the peptide and the extracellular matrix. In the biofilm control, macroscopic observation shows a homogenous whitish adhered layer covering the walls and bottoms of the well (Fig. 2D, middle), but when capsicumicine is present, we see whitish flocculent non-adhered heterogeneous agglutinates (Fig. 2*D*, right). Transmitted light microscopy (TLM) images match with these macroscopic observations (Fig. 2*E*). Crucially, SEM and TEM technique yield ultra-structural descriptions that support these results, with the control biofilm matrix showing denser assembled globular-like structures, while the capsicumicine-exposed matrix is less dense and has thin fibrillary branch-like structures (Fig. 2F and G). However, cellular morphologies are not different from the controls. Therefore, different imaging techniques prove that matrix assembly changes when capsicumicine is present.

### Capsicumicine interacts with matrix polysaccharides

To explore the peptide’s affinity with these major matrix components, we exposed *S. epidermidis* cultures to capsicumicine (-FITC; green) and saccharides staining, then analyzed them all by CFM. We used concanavalin A and calcofluor to selectively target matrix saccharides (blue). Amounts of capsicumicine-FITC appear exclusively on the matrix (Fig. 2*B* left; S2*B*) and its colocalization indicates the interaction between both marked elements (Fig. 2*C*, right; S2*A*). Visualization was done by individual (Fig. 2*C*, left and center) and colocalization fluorescence (Fig. 2*C*, right). The peptide control was pseudonajide-FITC, an antibacterial peptide; it showed green fluorescence in the cells but not in the matrix (Fig. 2*B* right; S2*C*). To explore whether capsicumicine features are compatible with carbohydrate-binding modules (CBMs), we performed a BLAST and amino acid alignments between capsicumicine and chitin-, chitosan-, and polysaccharide intercellular adhesin (PIA), PIA-binding proteins (Table S2). UniProt tools (*26*) and CAZy information crossing (*27*) showed that capsicumicine does in fact present CBM homology with all tested proteins (Fig. S3).

### Capsicumicine shifts Staphylococcal synthetic matrix

To confirm these interactions in the absence of bacteria metabolic or regulatory influences, we adapted a model of artificial staphylococcal biofilm assembly to test it. Briefly, we monitored the real-time molecular self-assembly (RTMSA) reaction by measuring the optical density at 600 nm (OD_600_) as a function of time with or without capsicumicine. OD increases when capsicumicine is present, which shows that the molecular self-assembly reaction is quicker overall (Fig. 3*A*). The profiles of assembled matrices are visually different, with larger agglutinates in the presence of capsicumicine, although for both controls are similar (Fig. 3*A*). Remarkably, these profiles are comparable to those previously observed in the presence of bacteria (Fig. 2*D*).

**Fig. 3.**
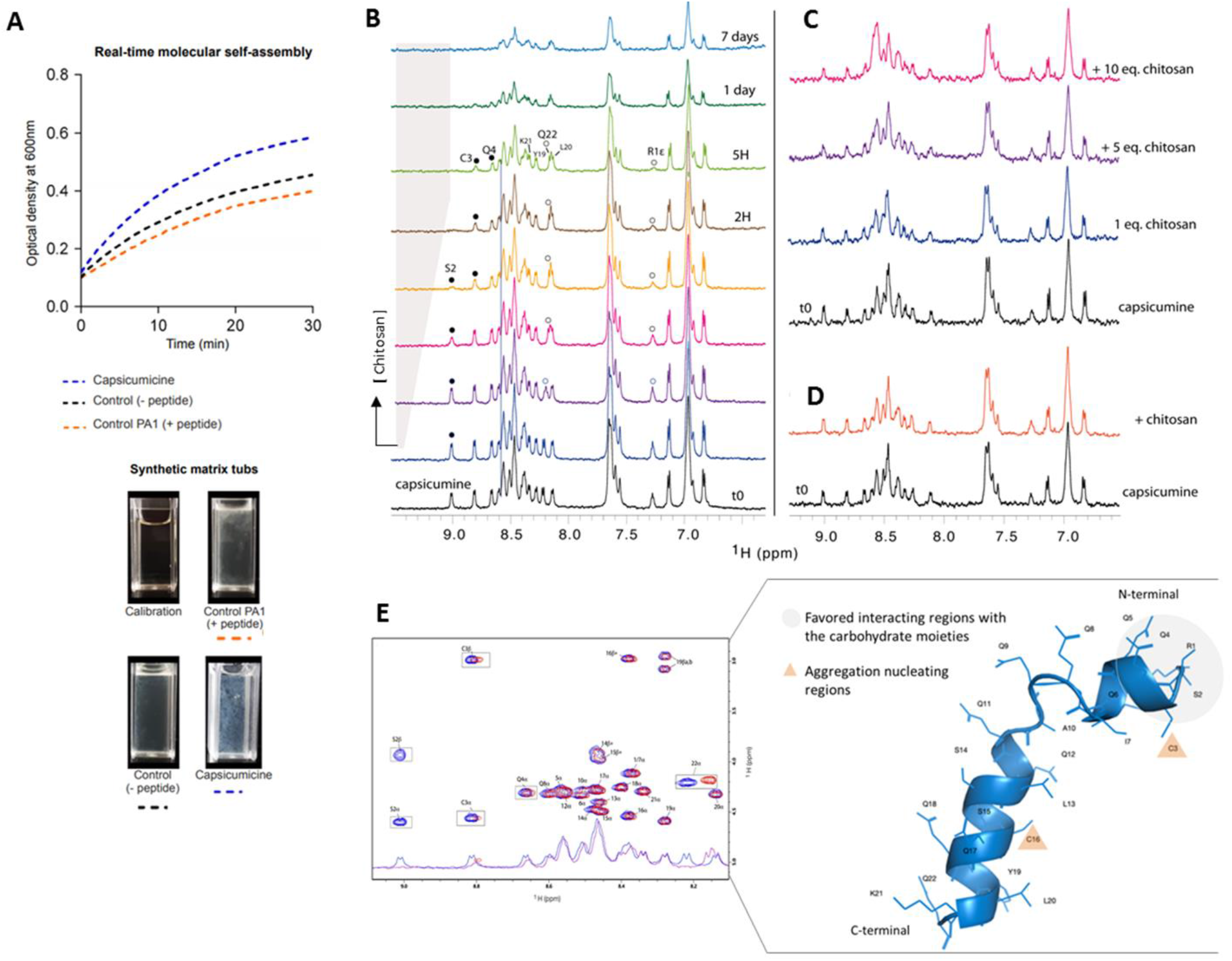
Molecular interactions between capsicumicine and target-saccharides. *(A)* RTMSA curves of artificial staphylococcal biofilm assembly. Optical densities (OD_600_) were recorded as a function of the time in the presence of capsicumicine (blue dots) or PA1 as peptide control (orange dots) and the synthetic matrix without peptides (black dots); each reaction tub is shown under the graph. *(B-D)* NMR titration of capsicumicine by chitosan at 600MHz, 10°C. *(B)* From the bottom to the top: increasing concentrations of soluble chitosan [< 0.5 eq. and 8% of volume variation] were gradually added to capsicumicine water solution. Specific line broadenings and frequency shifts are shown in the NMR spectrum for some amide protons upon titration; white bullets for terminal R1 and Q22 and black bullets for S2, C3 and Q4. *(C)* Assembled chitosan gradually added to soluble capsicumicine [0.5 eq.]; an overall line broadening was observed. *(D)* Assembled chitosan suddenly added to soluble capsicumicine [0.5 eq.]; the orange spectrum was recorded after few minutes of stirring. *(E)* Superposition of capsicumicine 1D NMR and TOCSY spectra immediately after sample preparation (blue) and chitosan [10 eq.] addition (red) and; capsicumicine structure predicted by Phyre2 server; according to NMR results the aggregation nucleating regions are shown as orange triangles and favored interacting region with sugar moieties is shown in grey.

To evidence the interactions between capsicumicine and target-saccharides we performed NMR titration experiments using chitosan as a mimic of the matrix poly-N-acetyl glucosamine (*PNAG*). The evolutions of the capsicumicine NMR spectra upon chitosan addition were monitored. Reference 1D proton NMR spectra were recorded at all tested conditions. Chitosan was gradually added from a concentrated solution and spectra changes were monitored. This solution remained clear during the 7 days of recording. The addition of soluble chitosan drives noticeable changes of the resonance frequency of some of the amide protons mainly in the N-terminal part of the peptide < S2, C3, Q4 and R1 (not shown) > (Fig. 3B). Conversely, to < Y19, L20 and K21 NH and the Y19 > aromatic proton resonances, between 6.8 and 7.2 are not sensitive to the presence of chitosan (Fig. 3*E*). The concentration variation of chitosan was estimated to less than 0.5eq at the end of the titration. This small quantity of chitosan drives spectral modifications in the N-terminal part of capsicumicine and strongly supports the interaction between both partners. The important broadening observed after 1 day shows that the peptide structure keeps evolving over time and this is not due to cysteines’ oxidation (Fig 4*D, E*). These experiments were repeated adding gelled chitosan instead of soluble chitosan, mimicking an assembled matrix. It does not drive any significant spectral changes, except for a general line broadening which probably arises from an increased viscosity of the solution (Fig. 3*C*). The suddenly addition of gelled chitosan pellets in the NMR solution, does not modify the spectrum either (Fig 3*D*).

### Structural requirements toward to molecular interactions with target-saccharides

To better understand the mechanisms involved in capsicumicine bioactivity we step toward the knowledge of its conformation in solution and molecular interactions with matrix representative saccharides. To monitor the conformation of free capsicumicine in response to time and temperature we first used Circular Dichroism (CD) and Nuclear Magnetic Resonance (NMR) spectroscopies. Spectral deconvolution using the CDSSTR algorithm discloses a helical structure (about 43-44%) with a non-negligible proportion of β strands (31-33%) and unfolded (about 20%) (Table S3). The structure stability was assessed over 5 days recording the CD spectra at 5°C (Fig. 4A). The main structure observed over the first 24h is helical. After 5 days, deconvolutions show a slight decrease of the helical content. Furthermore, on the fifth day, CD spectra were also recorded at 15 and 30°C (Fig. 4*B*). The helical content decreases to 18-23% while the β strand and turn mean proportions rose around 37% and 23%, respectively. The proportion of unfolded structures also increased. After 2 days at room temperature, a new CD spectrum was recorded at 30°C. The helix proportion became less than 15 % in favor of β strand and unfolded conformations (respectively 38-41% and 31-35%). During these days, the peptide solution remained clear, showing no macroscopic signs of aggregation.

The 1D and 2D NMR spectra of a 0.3mM freshly prepared solution of capsicumicine were recorded at 10°C, pH 5.0. All amide resonances were unambiguously assigned using TOCSY and NOESY spectra, except for Q9 and Q11 residues (Fig. S4). The spectral dispersion and the spreading of the amide proton resonances disclose that the peptide is folded with one conformation in these conditions. In accordance with CD spectra, a second set of NH resonances appears as a function of time (Fig. 4*C, D*), revealing conformational changes in the slow exchange regime on the NMR time scale. After 5 days, the most important chemical shift perturbations observed on the TOCSY spectrum are clustered around the two cysteines apart from the two terminal residues (boxed cross-peaks, Fig. 4*E*). Interestingly, I7 located one helix turn apart C3 is among the most sensitive residues to the conformational perturbation. This is due to a disulfide bridge (DB), since no reducing agent was added. Consequently, large proton chemical shift perturbations (CSP) would be expected for every - NH amino acid and this was not detected (Fig. 4*D*, *E*). The prevalence of the second conformation reaches more than 50% after 7 days (Fig. 4*C*, top). The solution remained clear overall the spectra recording time.

**Fig. 4.**
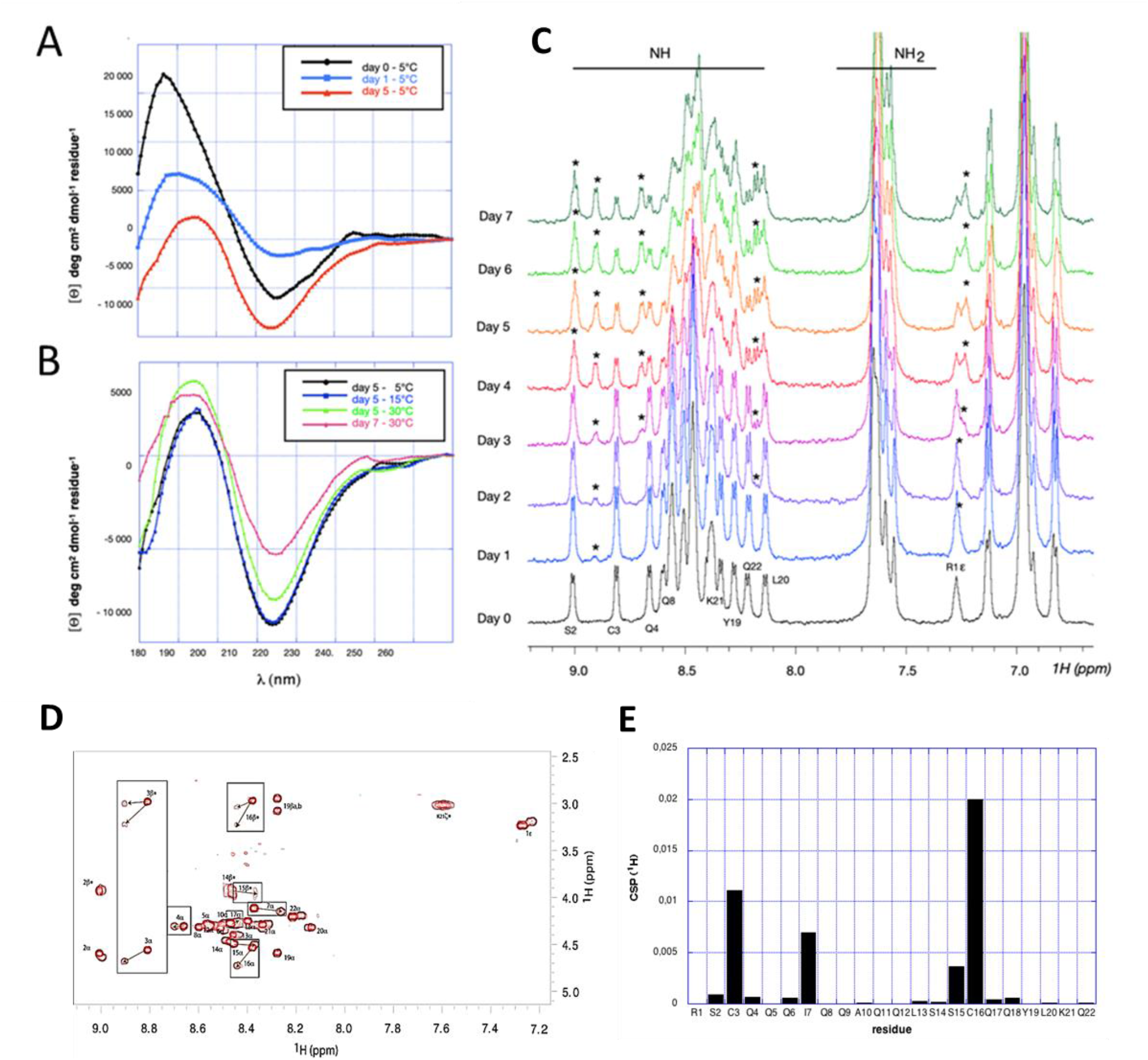
Capsicumicine conformational changes. CD spectras *(A)* Evolution at 5°C over 5 days; *(B)* Evolution *vs* time and temperature. *(C)* NMR spectra recorded at pH 5.0, 0.3mM and 10°C immediately after extemporaneous preparation (bottom); a second conformation appears over the time corresponding to the second set of amide resonances (stars). *(D)* Superposition of capsicumicine TOCSY spectra immediately after sample preparation (red) and at day 5 (brown). The most important chemical shit perturbations between the original and the new conformations are boxed and clustered around the two cysteines. *(E)* Proton chemical shift perturbations (CSP) computed from the TOSCY spectra are displayed in the box.

### Capsicumicine attenuates the dynamics of *S. aureus* (Xen36) infection on central venous catheter (CVC)

We performed a translational proof-of-concept evaluating long-term bacterial biofilm and related infection in mice implanted with capsicumicine pre-coated CVC. These CVCs were previously coated using an immobilization polymer (“hydrogel”) encompassing capsicumicine. This coating confers amorphous and biocompatible surfaces to CVC (Fig. 5*A*). They were first validated *in vitro*, decreasing ≥51 % of *S. aureus* colonization (Fig. 5*B*). Then, bacterial development was evaluated *in vivo* by bioluminescence imaging and bacterial load of harvested CVC (Fig. 5*C-F*). Two days (D2) after *S. aureus* systemic infection, 40% of the control group (CVC-Hydrogel) presented high bioluminescence signal (red zones) related to ROI against none in the treated group (CVC-Hydrogel + capsicumicine), (Fig. 5*C*). Four days (D4) after infection, 75% of the control presented high ROI red zones against 20% in the treated group (Fig. 5*C*). In addition, at D4, 1 animal was found dead in the control group. Therefore, bioluminescence quantifications show that the treated group decreased 56% (D2) and 54% (D4) of the total flux compared to the control (Fig. 5*D*). This trend was also observed on the CFU load at D4 (Fig. 5*E*). After image acquisitions (D4), 2 animals / group were euthanized due to ethical criteria. At the end of the experiment, 7 days after infection (D7), the treated group showed a decrease of 86% of bacterial load compared to control (Fig. 5*E*). Macroscopic observation revealed that one animal from the control presented several organs with necrosis (liver, spleen, intestine, kidneys and bladder). Finally, the treated group increased survival rate of 50% at D7 compared to control (Fig. 5*E*).

**Fig. 5.**
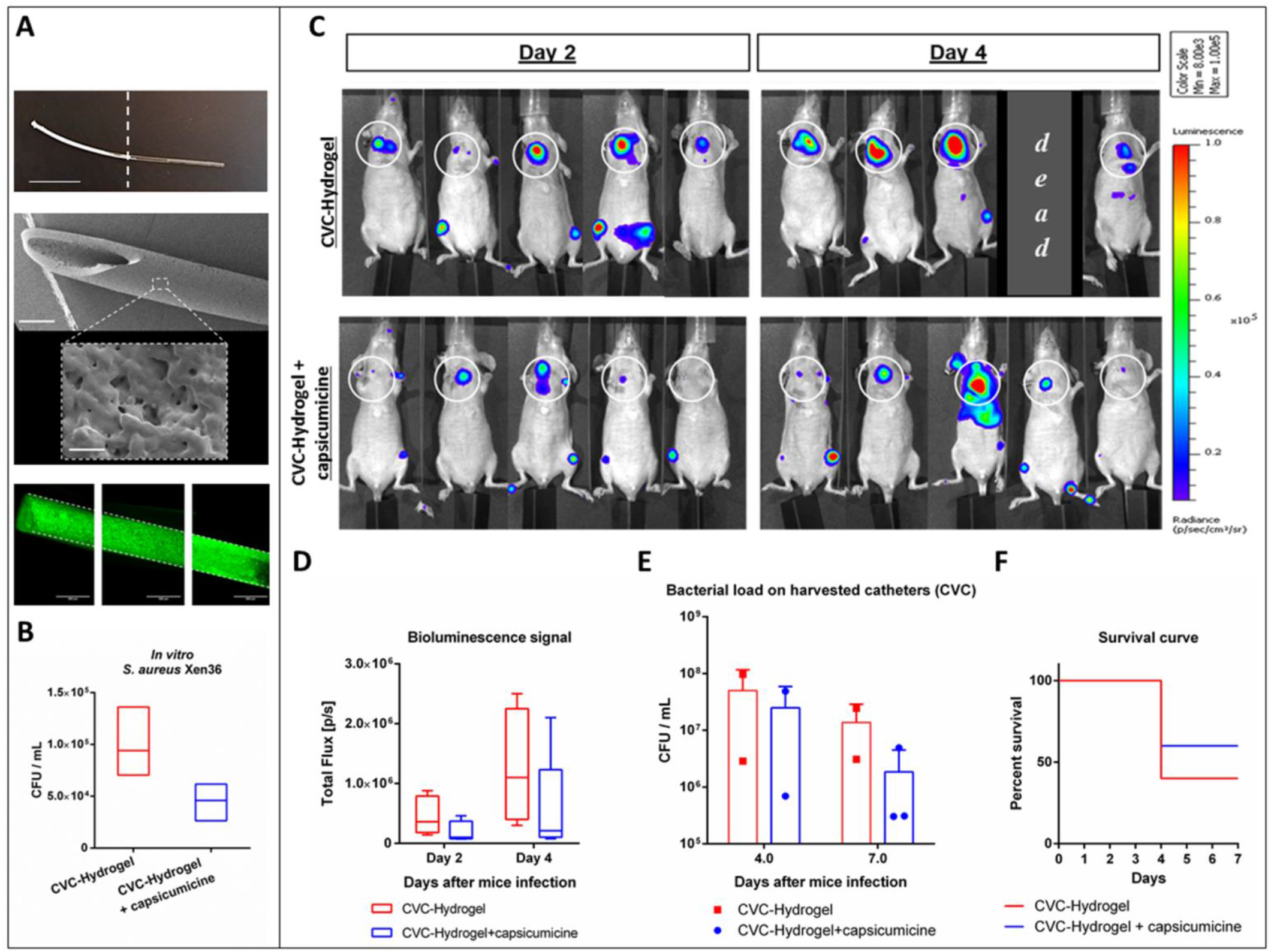
Assessment of capsicumicine pre-coated central venous catheter (CVC) in mice (SKH1) infected by *Staphylococcus aureus* Xen36. *(A-B) In vitro* CVC development; *(A)* From the top to the bottom: a picture of a up to half capsicumicine pre-coated CVC (whitish), scale bar 1 cm; SEM image of pre-coated CVC, scale bar 0.5 mm and its magnification, scale bar 1 μm and; a compilation of images of fluorescence microscopy of pre-coated CVC (capsicumicine-FITC in green); *(B)* Bacterial load on CVC. *(C-F) In vivo* CVC assessment; *(C)* IVIS images (radiance-thresholded and smoothed with a bidimensional Gaussian filter) of ventral position for each mouse at each time point; the control group (CVC-Hydrogel) is shown in the top and the treated (CVC-Hydrogel + capsicumicine) in the bottom; white circles delimitate the CVC localization; the luminescence scale is presented in radiance. *(D)* Bioluminescence quantification (ROI mean signals ± SD) is shown as photons total flux [p/s] for each group at each time point. *(E)* Bacterial load (CFU/mL) on harvested CVC for each mouse at each time point (day 4 and 7 after infection) and; *(F)* The Kaplan-Meier survival curve (% of survival) at each group, during 7 days after infection. All bioluminescence data and analysis were performed using IVIS Spectrum of Perkin Elmer. All experiments were conducted in accordance with ethical committee and the French authorities (n = 5 / group). Bioluminescence images are presented with a defined pseudo-color scale to visualize the intensity of signals emitted.

## Discussion

Bioinspired peptides are increasingly being explored as alternative biofilm controls, and become important allies in the fight against bacterial tolerance and resistance (*28, 29*). As shown here, capsicumicine, a peptide derived from *Capsicum baccatum* red pepper seeds, possesses strong antibiofilm activity *in vitro* and *in vivo*. Capsicumicine prevents the establishment and maintenance of biofilm architecture through a new mechanism of action the “matrix anti-assembly” (MAA). MAA differs from matrix *dis*assembly (*30*) as instead of de-structuring pre-established matrices it acts on the initial phase of assembly preventing its correct structuration. In fact, established biofilms are harder to treat than initial biofilms, because they have more complex structures (*31*), and increased structural complexity means that more energy is required for disassembly (*32*).

Bacterial surface proteins can passively interact with surfaces such as medical devices, generating an initial and reversible adhesion after electrostatic and hydrophobic interactions, Van der Waals forces and others (*33*). Bacteria will then require extracellular matrix production in order to remain attached after these weak interactions (*34*). During this process, physicochemical interactions drive molecular and colloidal matrix self-assembly, establishing a chain of dense architecture that results in stable adhesion (Fig. 6*A*) (*35*). In contrast, capsicumicine interacts with matrix saccharides and modifies the self-assembly chain, resulting in a less dense and nonfunctional matrix, and impaired biofilm formation (Fig. 6*B*).

**Fig. 6.**
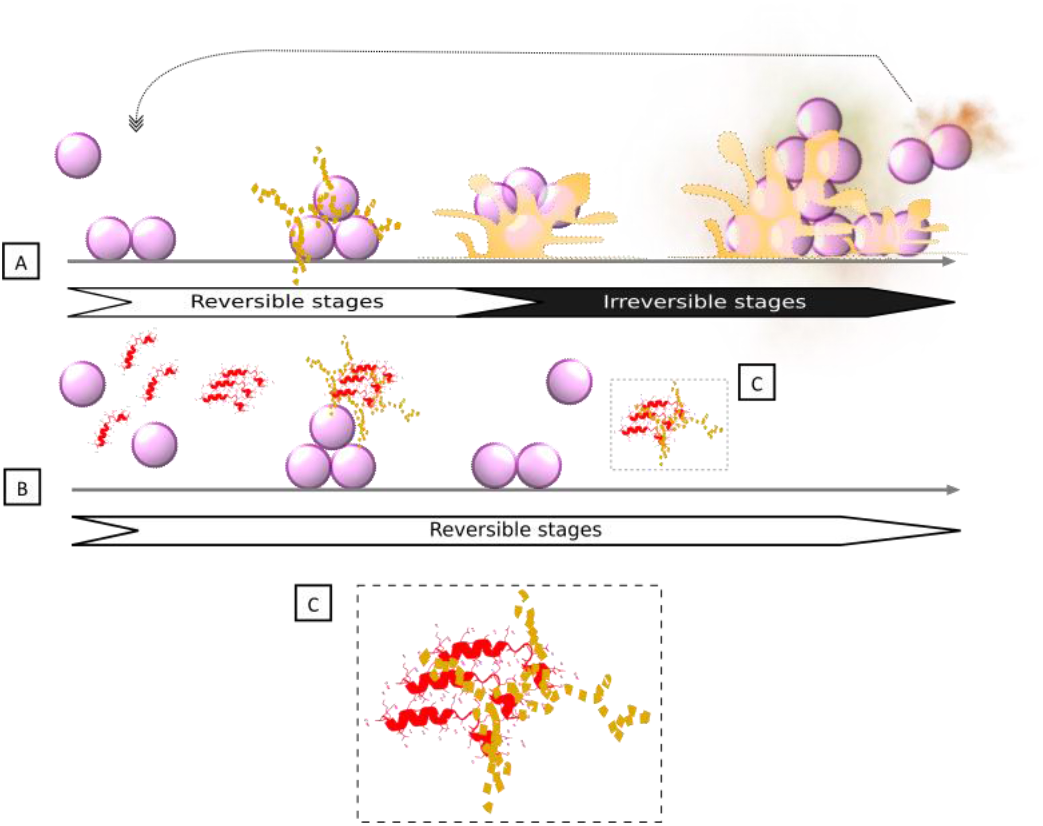
The matrix anti-assembly (MAA), antibiofilm mechanism of action (MoA) identified in capsicumicine. *(A)* Untreated biofilm development on abiotic surfaces: planktonic cells interact passively with the surface and start the extracellular matrix production. Initial adhesion is established, after which matrix self-assembly occurs, leading to irreversible biofilm structuring and adhesion. *(B)* Biofilm development in the presence of capsicumicine shows the MAA MoA: planktonic cells are still able to interact passively with the surface. At that point, capsicumicine acts, attracting extracellular saccharides and shifting the matrix self-assembly; *(C)* Capsicumicine (red) tends to organize itself in beta leaves favoring the interactions with matrix target-saccharides (yellow). These interactions occur between the non-catalytic carbohydrate binding modules of the peptide and the saccharides. This peptide supra organization can compete with matrix assembly, preventing/destroying intra matrix h-bonds over a large range of residues. Consequently, the MAA decreases adhesion and aggregation, forcing bacteria to remain planktonic. The arrows indicate reversible (white background) and irreversible stages (black background) during biofilm formation.

Capsicumicine’s amino acid sequence contains several residue characteristics for recognizing putative chitin-binding domains, including polar and hydrophobic residue (45% of Q) and cysteines (*36-39*). The chitin-binding domain is a well-conserved amino acid stretch that binds specifically to N-acetyl glucosamine, a homologous structure of PIA (*27, 40, 41*). Additionally, non-catalytic carbohydrate binding modules (CBMs) are contiguous amino acid sequences with a discreet fold displaying carbohydrate-binding activity. In this context, capsicumicine displays homologies with all tested CBM-proteins, notably with icaA protein that is a PIA synthetase from the same *S. epidermidis* strain studied here (Fig. S3 and Table S2).

According to the CAZY web site, there are 86 CBMs defined families up to now (http://www.cazy.org/Carbohydrate-Binding-Modules.html). CBMs exhibit different folds, with secondary structures ranging from mainly β-sheet based organization to a mixture of α-helices and β-sheets and usually display a high content of loops and unfolded parts. Likewise, capsicumicine secondary fold is a mixture of α-helix, β-sheet and unfolded regions, with conformational time and temperature interconversion in favor of β-sheet and unfolded segments. The β-sheet sub-structure observed in the CD concerns the Nter part of the peptide according to the chemical shifts of the corresponding amino acids (between 8.6 and 9.0). The TOCSY spectra analyses disclose that the conformational changes observed through the time are originated around the two cysteines, possibly involved in intermolecular disulfide bonds. This disulfide bond is possibly intermolecular since the formation of an intramolecular disulfide bond is only possible by bending the structure to get both cysteines close to each other. The size of different conformers and potential polymers are limited as shown by the NMR peaks of free peptides which are not broadened at day 7 (Fig. 4*C*). As a result, these conformational changes may correspond to structural requirements toward to interactions with staphylococcal matrix.

Antibiofilm, matrix anti-assembly (MAA) mechanism of action

This proposed mechanism is based on a set of intermolecular *(i)* and cooperative *(ii)* forces triggered by capsicumicine which then disrupt matrix assembly:

1. Intermolecular forces The staphylococcal matrix is composed mainly of polysaccharides, but also contains proteins (AtlE, Aap, Empb), teichoic acids, and extracellular DNA (*42*). These saccharides are PIA/PNAG, homoglycans of beta-1,6-linked 2-amino-2-deoxy-D-glucopyranosyl residues. The matrix contains positively-charged amino groups (PNAG), as well as negative charges from O-succinylation, which confers electrical charge lability on the matrix. Supported by NMR observations, capsicumicine interacts with free chitosan, a PNAG mimetic, and then switches to a higher molecular weight organization (Fig 3*B*). The line broadening observed with the peptide-chitosan mixture after one day evidence the aggregation of chitosan particles with capsicumicine over the time as schematized on Fig 6. Likewise, CSPs analysis reveals that chitosan is able to bind preferentially capsicumicine N-terminus (Fig 4*E*) before intermolecular disulfide bridges formation and the relative conformational transition. This is supported by the fact that chitosan is sub-stoichiometric at the end of the NMR titration and because the capsicumicine spectrum of the mixture is different from the spectra of the free peptide (Fig 4*C*). These interactions may proceed by both hydrogen-bonding between the hydrophilic Q’s lateral chains and hydrophobic/stacking interactions involving I and L residues. According to NMR, the N-terminal part of the peptide is probably the favored interacting region with the carbohydrate moieties. Thus, these data support the MAA model in which the peptide supra organization would compete with matrix self-assembly preventing intra h-bonds over a large range of residues. However, the NMR results obtained with assembled chitosan, mimetizing a preformed biofilm, demonstrate that capsicumicine does not interact with pre-assembled matrix (Fig 3*C,D*) such as demonstrated in the eradication test. Additionally, the amino acids in capsicumicine are mostly neutral, counting 13 Q and 3 S although, due to Q’s amide group and S’s hydroxyl group, they can generate electronegative (dipole-dipole) zones. This high electronegativity suggests that capsicumicine may interact with the positive free charges of the PIA/PNAG. Thus, these polar non-ionic forces perform moderate interactions with the polysaccharides. Since strong interactions such as ionic forces may trigger unwanted effects including matrix repulsion or sequestration (*43, 44*), the moderate interactions of capsicumicine seems to be ideal.
2. Cooperative polymerization forces In living systems, biomolecules perform their functions in the presence of various macromolecules of different shapes and sizes, and these interactions can include cooperative polymerization forces such as depletion forces (DFs) and subsequent molecular crowding (MC) (*45, 46*). These forces are noncovalent and non-specific physicochemical interactions leading to bridging, aggregation, and rheological variations (*47, 48*), as is observed in the presence of capsicumicine (Fig. *2D-G* and 3*A*), and also may contribute to MAA. In a suspension containing different molecules, DFs are the pressures exerted by small particles, which in turn cause attractive forces between the macromolecules. DFs are only expressed in crowded environments like biofilms, driving the assembly and final shape of these structures (*49*). Taking together NMR and CD experimental results as well as the TANGO and SALSA analysis (Fig. S5*B*), we can picture that capsicumicine can self-organize to form extended structures both by disulfide bridges and β-sheet mediated fibrils formation. We propose that capsicumicine-associated DFs may coagulate in polymer solutions forming fibers and parallel bundles (*50*), explaining the observed branch-like profile (Fig. *2D-G*) (*51*). This higher organization should facilitate the interaction with the saccharides of the matrix preventing their pattern assembly and resulting in flocculation (*52*). In a same way, MC in macromolecule solutions is characterized by a decrease in accessible volume due to high macromolecule concentrations, as well as attractive and repulsive forces between them (*45, 53*). Some molecular crowders modulate refolding kinetics and decrease competing aggregation and segregation (*54*). In this way, capsicumicine enhances the aggregation kinetics (Fig. 3*A*), acting as an agent of MC, decreasing entropic forces and leading to segregation (*55*). Thus, capsicumicine’ structures are chemically enable to interact with staphylococcal matrices saccharides. These combined intermolecular and cooperative forces disturb matrix self-assembly at both molecular and colloidal levels. Consequently, matrix functionality shifts and antibiofilm activity occurs. In other hand, planktonic microorganisms are more available for the innate immune system to recognize and clear them (*56*). Furthermore, non-antibiotic activity appears less susceptible to the development of bacteria resistance because these microorganisms are under less evolutionary pressure then when exposed to conventional antibiotics (*57, 58*). Finally, we demonstrated that capsicumicine strongly prevents biofilm formation for the most frequent bacteria related to nosocomial infections, *S. epidermidis* and *S. aureus*. We first confirmed that capsicumicine was not cytotoxic thus being safe for *in vivo* tests. Then, we trialed the peptide in a translational pre-clinical model that mimics medical device-related infection. As a result, pre-coated CVCs enhanced mice survival decreasing *S. aureus* colonization and consequently attenuated the infection. Although there are no antibiofilm drugs available here we relate the discovery of the first-in-class carbohydrate-binding peptide as a promising candidate for complementary drug/treatment of infectious diseases. In particular, we elucidated its antibiofilm mechanism of action, matrix anti-assembly (MAA) and validated a proof of concept towards to an *in vivo* application.

## Materials and Methods

### Peptides

All peptides were synthesized by Biomatik and ProteoGenix at purity grades over 95% in salts suitable for cell culture. For the assays, the peptides were all solubilized in ultra-pure sterile water.

### Bacterial strains and growth conditions

*Staphylococcus epidermidis* ATCC 35984 was grown overnight on blood agar (Thermo Scientific Oxoid PB5039A) at 37 °C. Oxoid LB agar was used for the colony-forming unit (CFU) assay. The other assays were done using a bacterial suspension of 3×10^8^ CFU/mL in tryptone soya broth (TSB, Oxoid) or 0.9% NaCl.

### Biofilm formation

At least three technical and biological replicates were done for each assay of 1, 10, or 100 μM peptide concentrations. *Biofilm formation inhibition:* A protocol adapted from Trentin et al (*59*) was used, with crystal violet in 96-well BD Falcon polyvinyl chloride (PVC) microtiter plates. The cell-bound stains were solubilized with absolute ethanol (Sigma-Aldrich), and absorbance was measured at 570 nm using a BIO-TEK PowerWave XS plate reader. The biofilm formation control represents 100% of biofilm formation. *Biofilm eradication:* Biofilm was pre-formed as described above for 24 h at 37 °C without treatment. Afterwards, the wells were washed to remove planktonic cells, peptide solutions and controls were added, and all were incubated for 24 h. Biofilm eradication was verified by evaluating the remaining content by crystal violet.

### Bacterial growth assays

*Microtiter plates:* Bacterial growth was evaluated by comparing OD_600_ values at the start and end of incubation in 96-well PVC microtiter plates. *Colony-forming units:* After incubation at 37 °C for 24 h, CFU/mL was calculated to determine the peptide solution’s bactericidal effects. The untreated growth control was considered to be 100% planktonic cells. At least three technical and biological replicates were performed for all assays.

### Quantitative reverse transcription PCR (qRT-PCR)

After culturing for 24 h, RNAs were isolated from planktonic controls, biofilm controls, and from total cells exposed to 10 μM capsicumicine. An Invitrogen TRIzol Max bacterial RNA isolation kit and an Ambion TURBO DNase treatment were used as per manufacturer instructions. Total RNA concentrations and purities were assessed using a Biochrom SimpliNano spectrophotometer, and PCR reactions was done to ensure the complete absence of DNA. Each qRT-PCR reaction was then subjected to previously established quantities of cDNA (10 ng) and primers (0.2 μM). Reactional volumes were calculated using SYBR Select Master Mix (Applied Biosystems), as per the manufacturer’s instructions. Primers (see Table S1) were designed using the Primer3 program, then produced by Eurofins Genomics. Applied Biosystems StepOnePlus equipment and software were used. Relative transcript levels were determined by the 2^-ΔΔct^ method (*60*).

### Cytotoxicity assays

Cytotoxicity assays were performed on the ImPACcell robotic platform (BIOSIT, Université de Rennes 1). Multiparameter high-content screening (HCS) and high-content analysis (HCA) were done on 7 different mammalian lines: HuH7, CaCo-2, MDA, HCT116, PC3, NCI-H727, and MCF7. The number of normal cells is presented as residual cell percentage compared to the DMSO control average.

### Microscopic analysis

*S. epidermidis* ATCC 35984 biofilm was cultured as described above.

### Scanning electron microscopy (SEM)

Sterile 10×4mm polystyrene coupons were inserted into bacterial cultures in the presence or absence of capsicumicine for 1, 4, and 24 h. The coupons were then washed with sterile 0.9% NaCl and fixed with 2.5% glutaraldehyde, 2% paraformaldehyde, and 0.1 M cacodylate buffer (pH 7.2). Afterwards, they were washed with 0.1 M cacodylate buffer and 0.2 M sucrose, then dehydrated with increasing concentrations of ethanol. A Leica EM CPD300 was used for critical point drying of the dehydrated samples. These were then sputtered with palladium in a Leica EM ACE200, and analyzed with a JEOL JSM-7100F microscope with EDS and EBSD at 10 kV.

### Transmission electron microscopy (TEM)

All well content was carefully detached at 1, 4, and 24 h, centrifuged at 10,000 g for 15 min at 4 °C, then washed with sterile 0.9% NaCl. Fixation was performed at 4 °C with sodium 0.1 M cacodylate, 2% paraformaldehyde, 2.5% glutaraldehyde, and 75 mM lysine. Samples were washed with 0.1 M sodium cacodylate and 0.2 M sucrose and contrasted with 1% osmium tetroxide and 1.5% potassium ferrocyanide. Dehydration was done with a gradual solution of ethanol and infiltration of increasing concentrations of LR White resin (Delta Microscopies, France). LR White resin inclusion and polymerization were then performed over 24 h at 60 °C in the absence of O_2_. Thin 80 nm sections were collected onto carbon grids, and visualized at 200 kV with a FEI Tecnai Sphera microscope equipped with a Gatan 4k x 4k CCD UltraScan camera.

### Confocal fluorescence microscopy (CFM)

Capsicumicine-fluorescein isothiocyanate (capsicumicine-ITC, 10 μM) was used to detect the capsicumicine peptide, whose antibiofilm activity was previously verified. After incubation for 1, 4, or 24 h, the well contents were carefully detached, centrifuged at 11,000g for 2 min at 4 °C, then washed with sterile 0.9% NaCl. The suspension was visualized directly or after adding 0.1μg/μL concanavalin A conjugates (Alexa Fluor® 633, Invitrogen) or 2 mg/mL Calcofluor White dye (Fluorescent Brightener 28, Sigma-Aldrich). To find bacterial cells permeated by the peptide control, we used pseudonajide FITC-labelled antimicrobial peptide (Ref). Images were acquired via resonant scanner with a Leica SP8 DMI 6000 CS confocal microscope with hybrid detector, and ImageJ software was used for image analysis.

### Real-time molecular self-assembly (RTMSA) assay

After checking the starting point (pH 7.2) of the assembly reaction for the synthetic staphylococcal matrix (*35*), we recorded the OD_600_ as a function of the time every 30 sec until 30 min. Molecular self-assembly reactions were calculated to a final volume of 4 mL, with 0.3% chitosan (medium molecular weight, 75-85% deacetylation), 0.15% bovine serum albumin, and 0.015% lambda DNA (all from Sigma) in TSB. The concentration (μM) of tested peptides was calculated for a final volume of 4 mL. Before getting the assembly reaction pH starting points, a calibration record was done using the same reactional tube containing all reagents (auto zero). Acetic acid and NaOH were used to adjust pH, and the reaction temperature was about 30 °C. As a negative control a similarly sized peptide was used, PA-1 (*61*).

### NMR

NMR spectra were recorded in 3mm tubes on a Bruker Avance III 600MHz spectrometer equipped with a TXI (1H,13C,15N) probe and a Z-gradient unit. Spectra were processed with Topspin 4.0.8 (Bruker Biospin) and CcpNmr Analysis (*62*). TOCSY and NOESY experiments were respectively recorded at 10°C and pH5.0 with a 70ms and 300ms mixing time. Capsicumicine concentration in water was set at 0.3mM and the pH adjusted either 5.0 or to 3.5 with few microliters of deuterated HCl 0.1M and/or NaOH 0.1M solutions. Chitosan powder (purchased from Sigma) was dissolved in pure water and the pH adjusted to either 5.0 or 3.5. Reference 1D proton NMR spectra were recorded at 10°C and either at pH3.5 and 5.0. Chitosan was stepwise added from a concentrated solution at pH 5.0 or 3.5 and spectra changes were monitored. We first titrated capsicumicine solution with chitosan solution at pH 5.0. The concentration variation of chitosan was estimated using the peaks intensities, resulting in 3.6 and 3.8 ppm (not shown) in the beginning to less than 0.5eq at the end of the titration. These experiments were repeated with a starting capsicumicine solution at pH 3.5 and the gradually addition of gelled chitosan, at pH 3.5. The suddenly addition of gelled chitosan pellets in the NMR solution, at pH3.5.

### Circular Dichroism (CD)

The peptide was dissolved in 18M⍰ water at a concentration of 50μM, at pH 5.0. The ultraviolet CD spectra were recorded at 5, 15 and 30°C, in a 0.1cm path length quartz cell on a Jobin Yvon CD6 spectrometer equipped with a temperature controller unit, over a 180-260 nm range, with a 2nm bandwidth, a step size of 1 nm and an integration time of 2s per point. The samples were conserved at 5°C between each recording. Spectra were averaged over 5 records. Water CD contributions were subtracted from CD spectra before processing. Spectra were processed using Kaleidagraph (Synergy Software). Molar circular dichroism (⍰⍰) per residue and molar ellipticity per residue (⍰⍰], MER) were computed from the difference of the delta absorbance recorded by the spectrometer. Raw delta epsilon per residue spectra were analyzed using the CDSSTR program (*63*) and different reference protein data sets provided by the DICHROWEB facility (*64*). MER data curves were smoothed for presentation, using the interpolate and weighted data (5%) routines provide by Kaleidagraph.

### TANGO (*65*) and SALSA (*66*) predictions

Tango predictions were run at 298K. Predictions were identical for pH set either to 5.0, 7.0 or 9.0. SALSA predictions were run with a window size dynamic of 4-20 residues, a cutoff of 1.2 and a minimal hot spot length of 5.

### CVC coatings

We adapted an approach to immobilize peptides based on poly(ethylene glycol)diacrylate (PEGDA) hydrogel (*67*). Briefly, polyurethane (PU) tubes (Instech) of 2Fr or 25G equivalent were used as coating framework. Hydrogel base was prepared using pentaerythritol tetrakis(3-mercaptopropionate) - PTMP (4.1 mmol), PEGDA (10 mmol), PEG-600 (20mM), 2,2-Dimethoxy-2-phenylacetophenone - DMPA (0.1 wt%), THF soluble PU (10 wt%) and methanol (qs). Peptides were solubilized in DMSO and added to hydrogel base under vortex agitation. PU tubes were first internally coated by suction using a needle-syringe and then externally by immersion. The reaction and polymerization conditions such as room temperature and oxygen tolerance were convenient. After polymerization, successive washes with methanol under agitation were used to eliminate undesirable monomers. All reagents were purchased from Sigma Aldrich.

### CVC infection assay

This study was conducted by Voxcan s.a.r.l. The ethical agreement is part of the project n° APAFiS# 10756-2017072522272676 v4, approved by Voxcan ethical committee (CEAA-129) and the French authorities (ministry of national education, the higher education and research). This study used SKH1 mice, females, immunocompetent and specific pathogen free (SPF) provided by Charles River Laboratories. Animals were acclimated at least 2 days, housed collectively in disposable standard cages in ventilated racks A3, at +21 ± 3°C, 30-70% of humidity, 12 hours of dark and light cycles, with filtered water and autoclaved standard food provided *ad libitum*. Catheters were blind implanted (n = 5 / group) in mice jugular vein following by an intravenous (IV) inoculation of *S. aureus* Xen36, 5 days days later, which was in turn to colonize the device from the blood circulation. The bacterial development was evaluated and compared between the different catheters by *in vivo* bioluminescence imaging (IVIS Spectrum, acquisition and analysis with Living Image 4.5.5 version) performed 2 and 4 days after mice infection. In addition, 7 days after inoculation or at the time of mice euthanasia for ethical reason, catheters were harvested and bacterial load were evaluated by CFU counting (SCAN500). All along the study, mice clinical state was evaluated using a scoring grid and body weight measurements 3 times a week. At each timepoint, a bioluminescence acquisition was also performed on the background (BKG) mouse to measure the flux level corresponding to the auto-bioluminescence. Images were radiance-thresholded with respect to the background radiance level and smoothed with a bidimensional Gaussian filter (3×3).

## General

We thank Daniel Thomas for his support, and Juliana Berland for insightful comments on the manuscript. Thanks to the Microscopy Rennes Imaging Center (MRic) and CMEBA facilities.

## Author contributions

R.G.V.B, S.C.B.G, A.J.M, S.N-L. and R.G. designed research; R.G.V.B, S.C., R.S., S.N-L. and S.B. performed research; R.G.V.B, A.R.Z., E.G., S.C.B.G, A.J.M, S.N-L., S.B. and R.G. analyzed data; and R.G.V.B, A.J.M, S.N-L. and R.G. wrote the paper.

## Data and materials availability

All data needed to evaluate the conclusions in the paper are present in the paper and/or the Supplementary Materials. Additional data related to this paper may be requested from the authors.

## Supplementary Materials

**Table S1.**
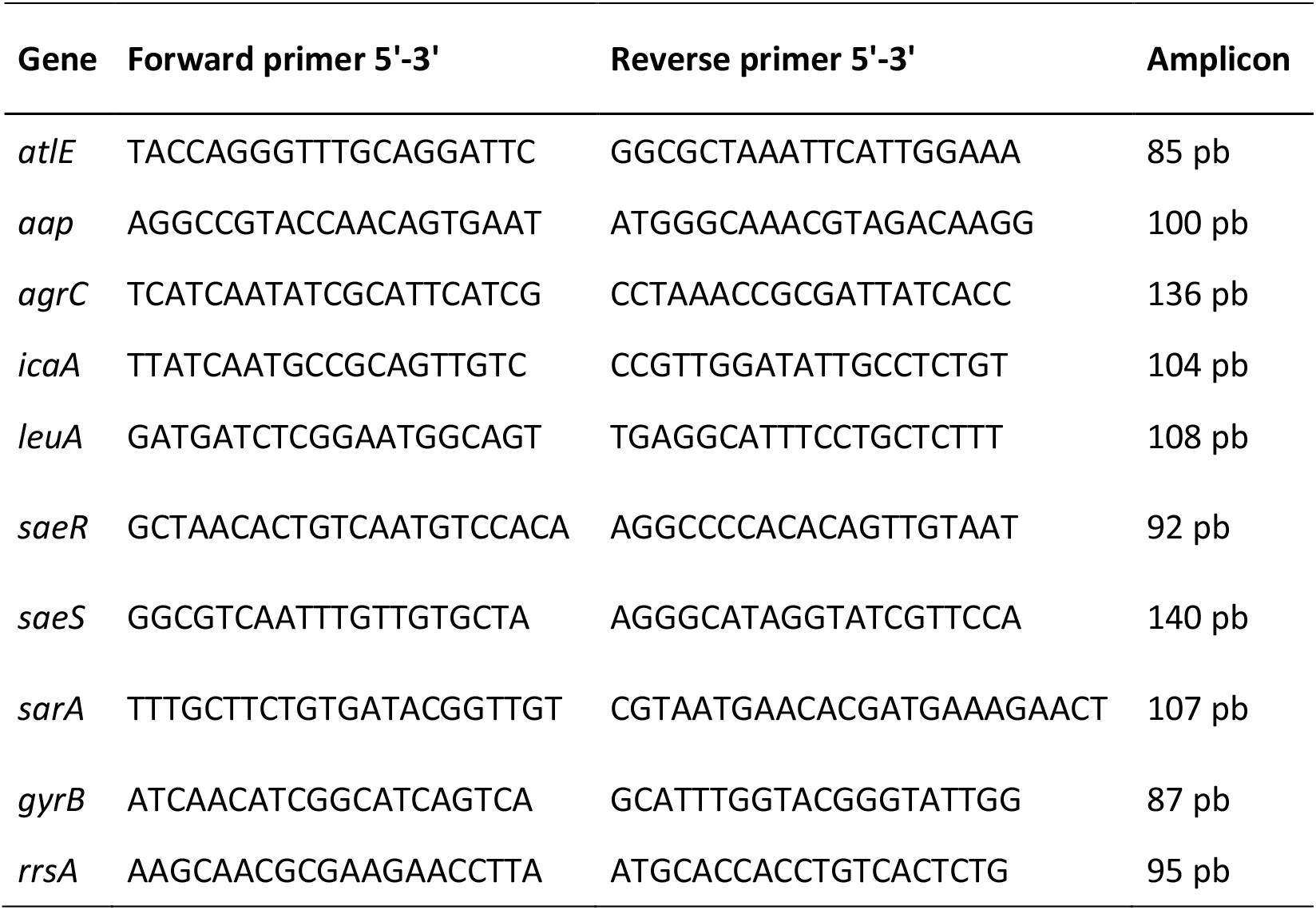
Primers used in this study. These were previously designed using the Primer3 program and standardized by our team.

**Table S2.**
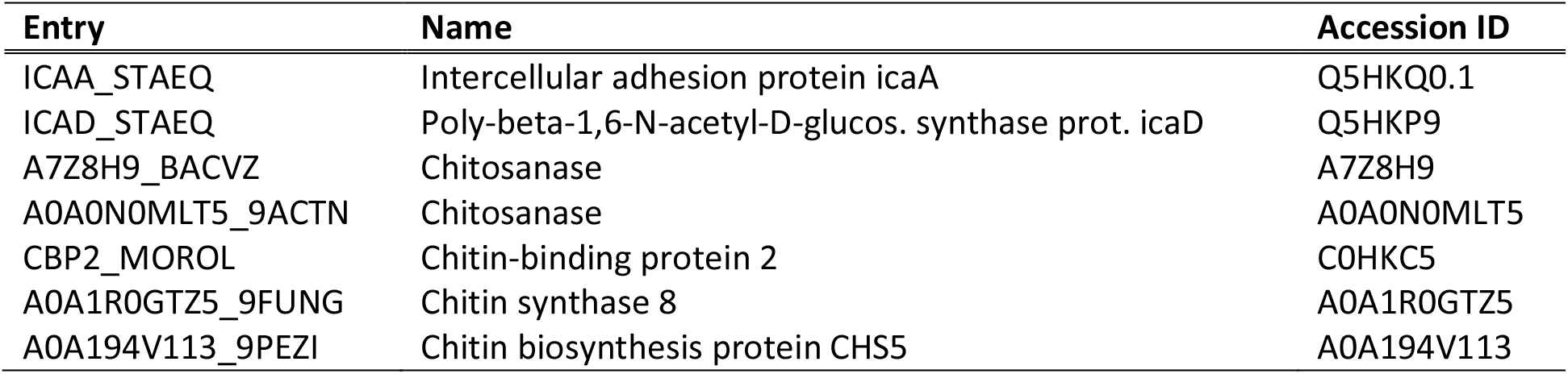
Carbohydrate-binding domain proteins that are capsicumicine homologs. These proteins are available in the UniProtKB database and were used for BLAST and amino acid alignments to perform similarity analysis.

**Table S3.** Results of the CD spectra deconvolution using CDSSTR algorithm.

| Day | Temp (°C) | Helix (%) | Bêta (%) | Turn (%) | Unfolded (%) |
| --- | --- | --- | --- | --- | --- |
| 0 | 5 | 42-46 | 29-34 | 5-8 | 16-21 |
| 1 | 5 | 42-45 | 30-32 | 6-8 | 20-21 |
| 5 | 5 | 38-42 | 24-31 | 8-12 | 17-23 |
| 5 | 15 | 22-26 | 36-45 | 13-14 | 20-26 |
| 5 | 30 | 16-24 | 33-40 | 18-20 | 20-25 |
| 7 | 30 | 6-17 | 38-41 | 10-23 | 31-35 |

**Fig. S1.**
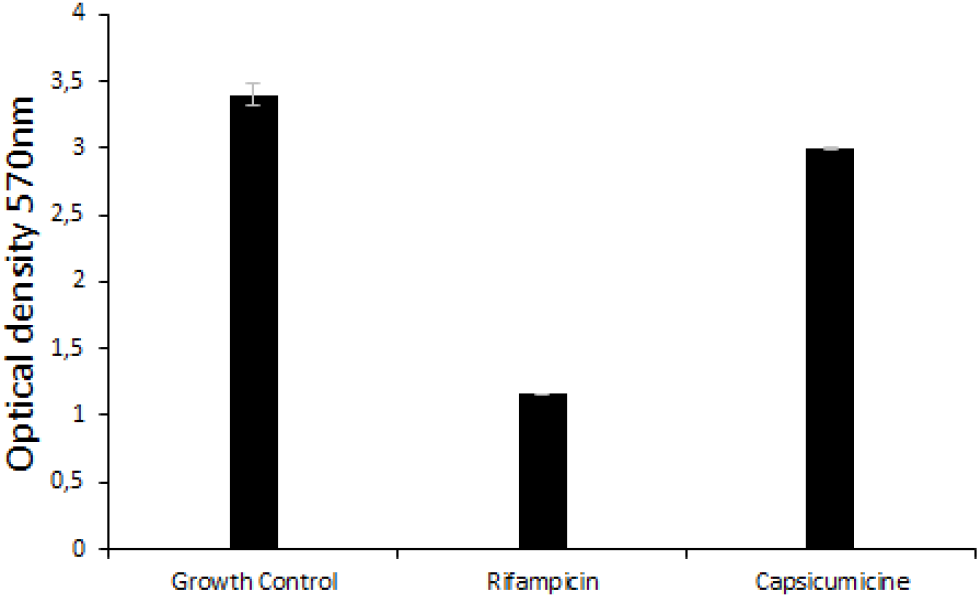
Biofilm eradication test. Shown are *Staphylococcus epidermidis* (ATCC 35984) biofilm quantifications at OD_570_ for the bacterial biofilm without peptide exposure (“Growth Control”), after exposure to the rifampicin antibiotic control, and after 24 h treatment with 100 μM capsicumicine.

**Fig. S2.**
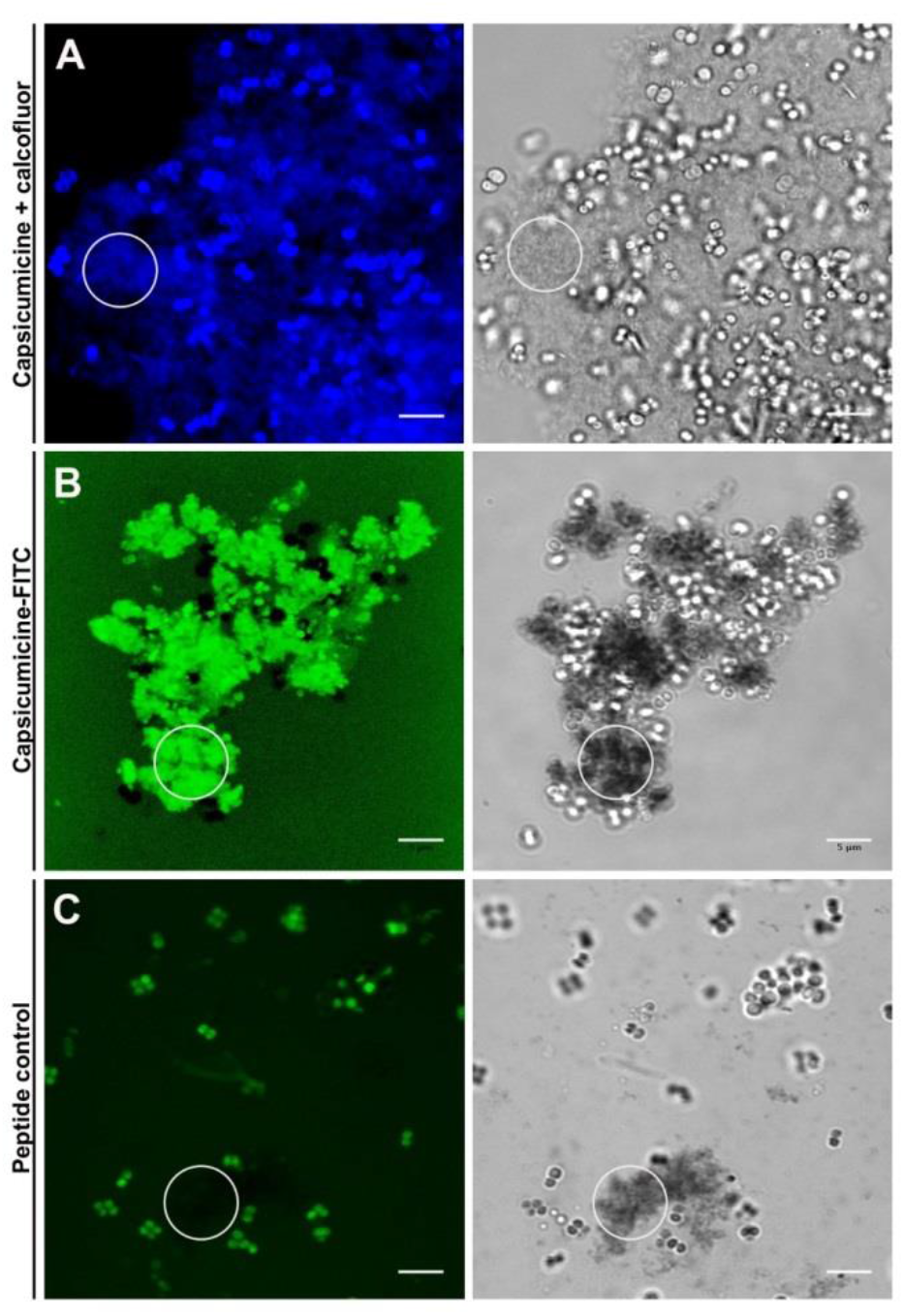
Confocal fluorescence microscopy (CFM) of *Staphylococcus epidermidis* (ATCC 35984). Calcofluor was used to highlight matrix polysaccharides (blue), and FITC used for the peptides (green). Visualization was done by fluorescence (left) and transmitted light (right) microscopy. (*A*) Cultures exposed to capsicumicine and calcofluor. (*B*) Cultures exposed to capsicumicine-FITC. (*C*) Peptide negative control cultures exposed to an antimicrobial peptide-FITC (Pseudonajide). The matrix is highlighted (circles). Scale bars, 5 μm.

**Fig. S3.**
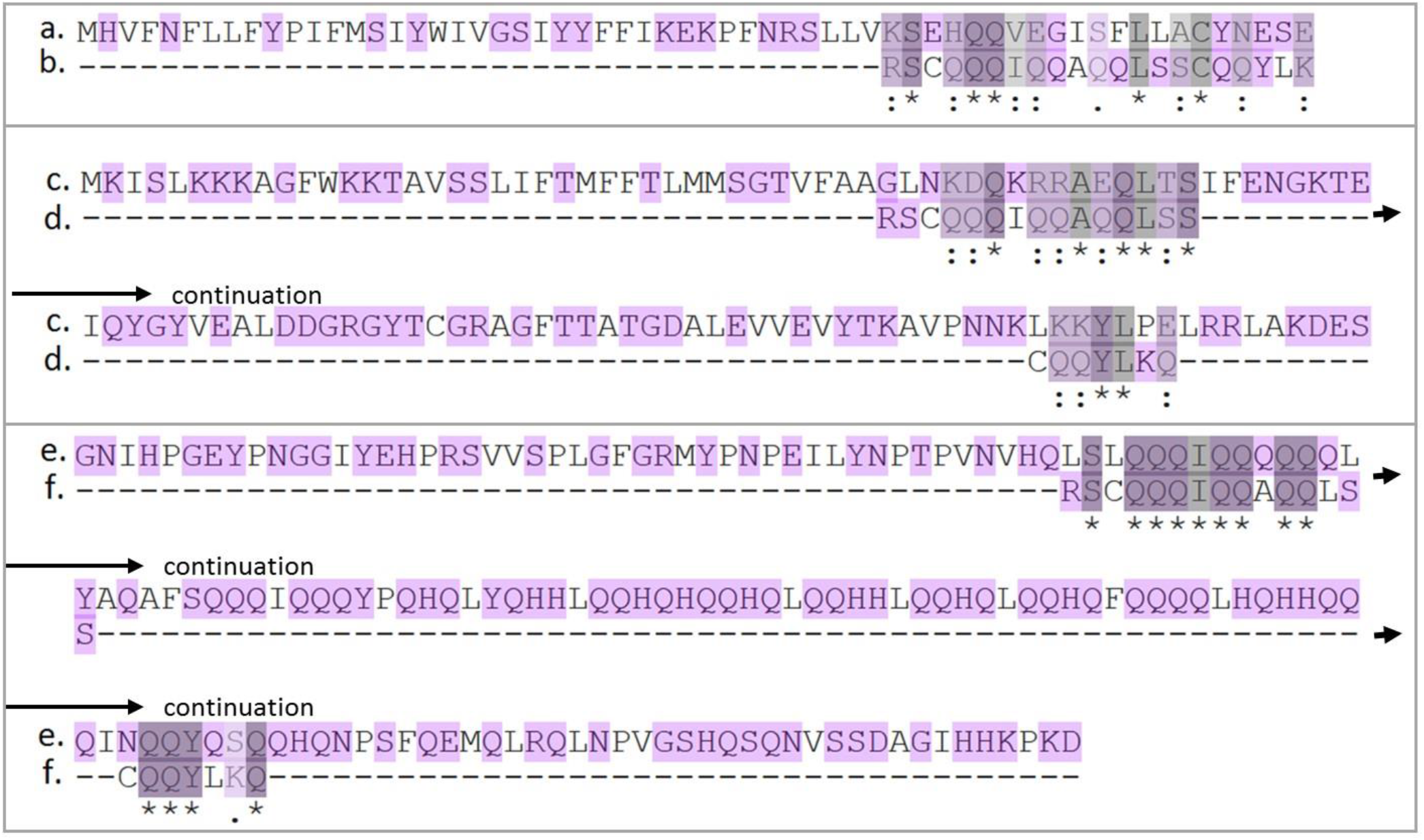
Amino acid alignment of capsicumicine and its carbohydrate-binding domain protein homologs. (*a*) The icaA (protein accession number Q5HKQ0.1) fragment from 1 to 60. (*b*) Capsicumicine fragment from 1 to 21. (*c*) Chitosanase (A7Z8H9) fragment from 1 to 60 and from 61 to 120. (*d*) Capsicumicine fragment from 1 to 15 and from 16 to 22. (*e*) Chitin synthase 8 (A0A1R0GTZ5) fragment from 1801 to 1860 and from 1921 to 1967. *(f)* Capsicumicine fragment from 1 to 14 and from 16 to 22. Equal (* and grey highlighting), similar (.), and highly similar (:) amino acids are indicated, as well as amino acid polar characteristics (purple). Support for this analysis is available at UniProt.

**Fig. S4.**
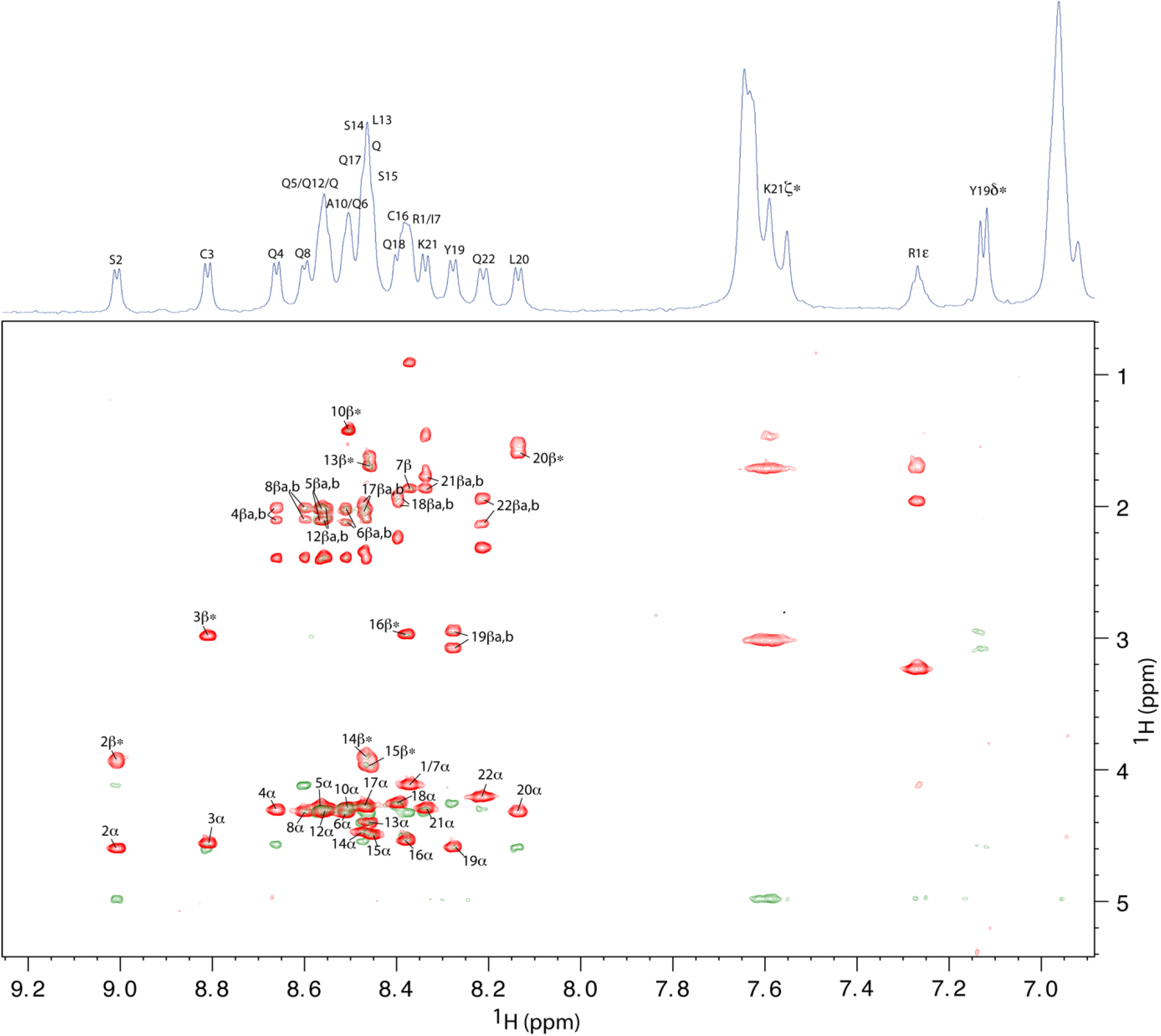
Superposition of the TOCSY (red), NOESY (green) and 1D (blue) spectra of capsicumicine recorded at pH 5.0 and 10°C. All amide NH resonances were assigned using nNH-(n-1)Hα NOESY cross-peaks, except for Q9 and Q11 residues due to poor resolution. For clarity, only NH-Hα and NH-Hβ labels were reported. The specta display 22 spin systems as expected for a single conformation. According to the spectral dispersion, capsicumicine is mainly folded.

**Fig. S5.**
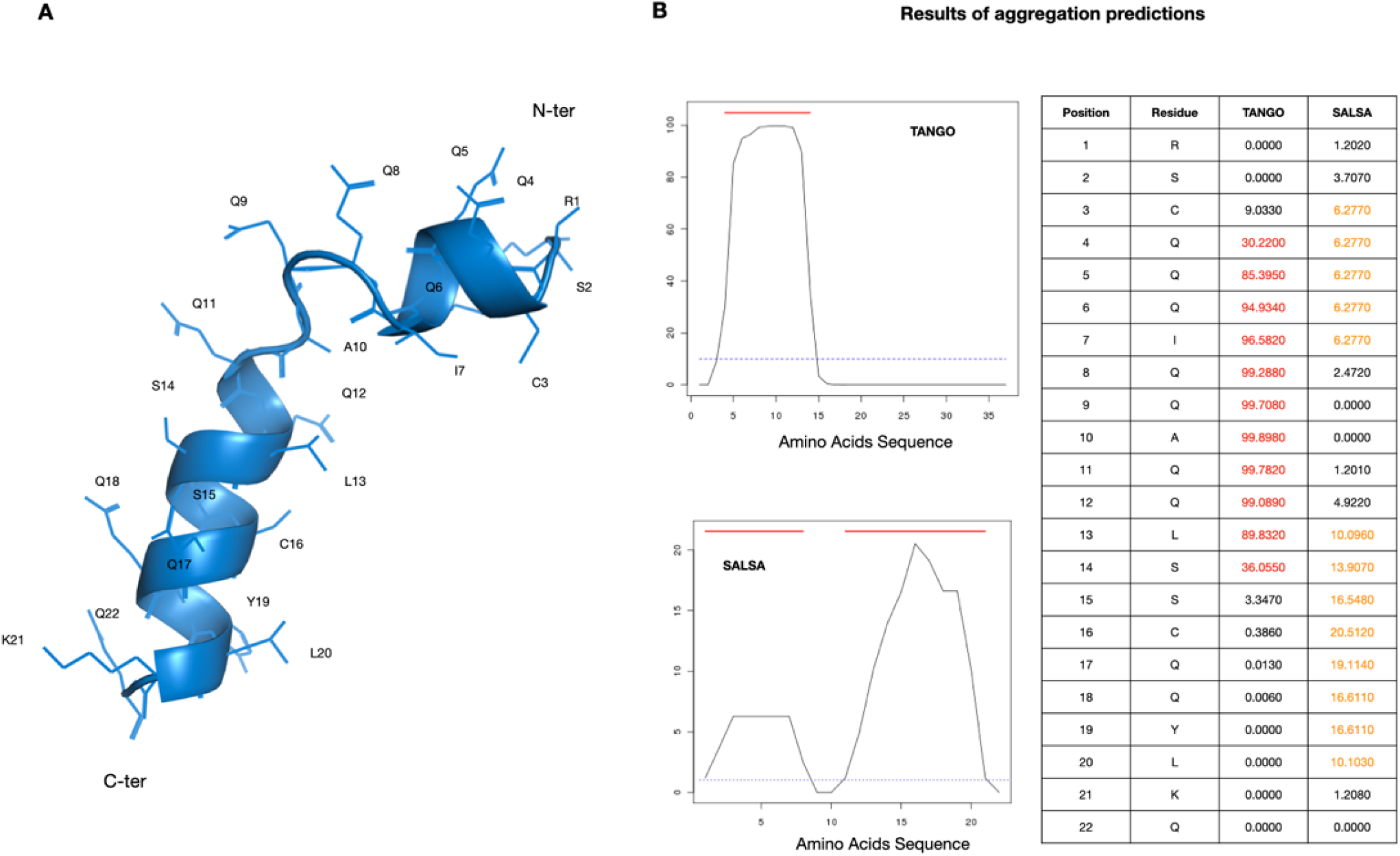
(***A***) Predicted capsicumicine structure using the Phyre^2^ prediction server. (***B***) Aggregation structure predictions using TANGO (*68*) and SALSA algorithms (*66*).

